# The homeodomain drives favorable DNA binding energetics of prostate cancer target ONECUT2

**DOI:** 10.1101/2023.06.13.544830

**Authors:** Avradip Chatterjee, Brad Gallent, Madhusudhanarao Katiki, Chen Qian, Matthew R. Harter, Michael R. Freeman, Ramachandran Murali

**Affiliations:** Department of Biomedical Sciences, Research Division of Immunology, Cedars-Sinai Medical Center, Los Angeles, CA 90048. USA; Samuel Oschin Comprehensive Cancer Institute, Cedars-Sinai Medical Center, Los Angeles, CA 90048. USA; Department of Urology, Cedars-Sinai Medical Center, Los Angeles, CA 90048. USA

## Abstract

The ONECUT transcription factors feature a CUT and a homeodomain, evolutionarily conserved elements that bind DNA cooperatively, but the process remains mechanistically enigmatic. Using an integrative DNA binding analysis of ONECUT2, a driver of aggressive prostate cancer, we show that the homeodomain energetically stabilizes the ONECUT2-DNA complex through allosteric modulation of CUT. Further, evolutionarily conserved base-interactions in both the CUT and homeodomain are necessary for the favorable thermodynamics. We have identified a novel arginine pair unique to the ONECUT family homeodomain that can adapt to DNA sequence variations. Base interactions in general, including by this arginine pair, are critical for optimal DNA binding and transcription in a prostate cancer model. These findings provide fundamental insights into DNA binding by CUT-homeodomain proteins with potential therapeutic implications.

**One-Sentence Summary:** Base-specific interactions regulate homeodomain-mediated stabilization of DNA binding by the ONECUT2 transcription factor.

## Introduction

The ONECUT (OC) transcription factors, consisting of OC1, OC2 and OC3, are associated with development of gastrointestinal organs (*1-7*) as well as neuronal components including the retina (*8-10*) and motor neurons (*11*). Recent studies have identified OC2 as a master transcription regulator driving lethal and therapy resistant prostate cancer (PC) (*12, 13*). In metastatic PC, OC2 is upregulated, promoting treatment resistance and transdifferentiation to neuroendocrine PC (NEPC) through repressing the androgen receptor (AR) axis, and activation of *PEG10*, a known NE driver (*12*). In addition, OC2 overexpression can also promote NEPC development by regulating hypoxia signaling (*13*). Furthermore, an OC2 inhibitor suppressed tumor growth and metastasis in a xenograft mouse model. OC2 has thus emerged as an important cancer therapeutic target, especially in treatment-resistant prostate cancer. Therefore, a better molecular understanding of this transcription factor is of fundamental and therapeutic importance.

The OC family proteins feature a conserved DNA binding module comprised of a CUT domain and a homeodomain (HOX) that are separated by a flexible linker, an arrangement also seen in POU transcription factors (*3, 8, 14-16*). CUT, like the structurally homologous POU-specific domain, shares a similar fold as λ and 434 phage repressor DNA binding motifs (*17-20*) while HOX is a widespread gene regulatory element with nearly 30% representation in transcription factors in humans (*21*). Thus, CUT and HOX represent two of the most evolutionarily conserved DNA binding elements essential in development. Prior structural work on POU members (*19, 22*) and subsequently OC1 (*23*), show the CUT and HOX bound to the major groove of DNA in an ‘overlapping’ manner to the opposite strands. However, a detailed analysis of the mechanism of DNA binding of CUT-HOX proteins is lacking. As a result, the specific roles of the CUT and HOX domains, as well as insights into the determinants of cooperativity, remain poorly resolved.

To obtain mechanistic details into these processes, and in the context of establishment of OC2 as a key player in PC, we carried out structural characterization of the OC2 DNA-binding module (OC2 hereafter) in complex with a *PEG10* promoter DNA sequence. We pursued these structural observations to carry out a stepwise biophysical analysis to understand the DNA binding by OC2 while further validating the results in a prostate cancer model.

## Results

### Structure of human OC2 in complex with *PEG10* promoter (*PEG10*) DNA

The OC2-*PEG10* structure resembles that of a previous OC1-*TTR* complex (*23*) (Fig. 1A and fig. S1A). Like OC1, the two α-helical domains of OC2, CUT and HOX, together with the connecting linker, wrap around the DNA major groove while absence of electron density of the linker residues suggests it does not physically bind the DNA. The CUT domain (amino acids 330-407) forms five alpha helices (α1-α5) while the HOX domain (amino acids 427-481), positioned at the C-terminal of CUT, forms three α-helices (α6-α8). The helices α3 of CUT and α8 of HOX each insert into the DNA major groove (Fig. 1, B and C). Compared to HOX, CUT makes most contacts with the DNA, a significant number of which are DNA backbone mediated (Fig. 1D). As a result of these contacts, the *PEG10* DNA is mildly distorted, with a slight bulge in the major groove region, compared to the canonical B-DNA (fig. S1B).

**Fig. 1.**
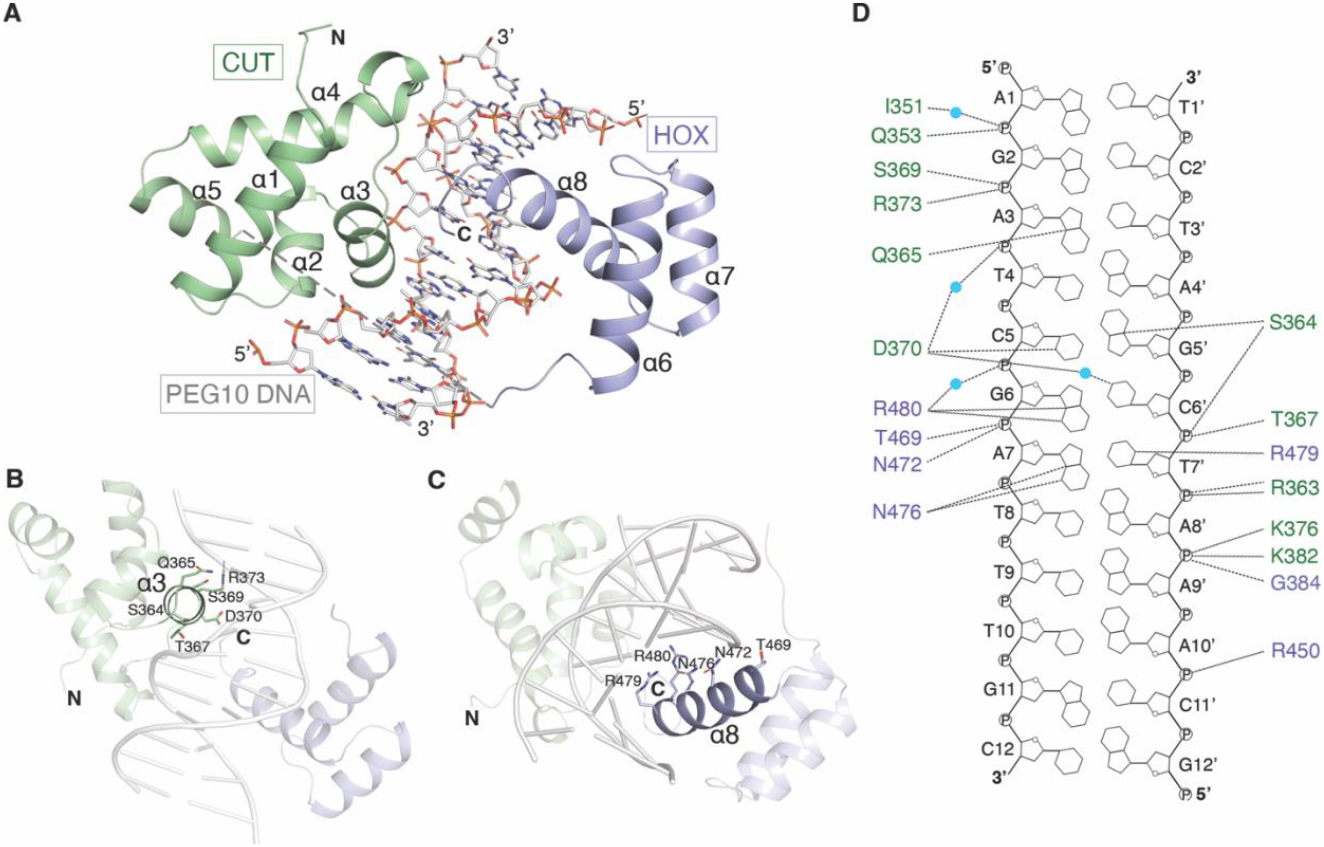
Structure of the OC2-*PEG10* complex. (**A**) Overall structure of the OC2-*PEG10* DNA complex. The position of CUT and HOX domains on DNA and their respective helices are labeled (α1-α8) while the unmodelled loop is depicted as a dashed line. CUT and HOX domains are shown in green and blue, respectively. Arrangement of DNA interacting residues in (**B**) α3 helix of CUT domain and (**C**) α8 helix of HOX domain of OC2 are shown. (**D**) Schematic representation of the protein-DNA contacts in the complex. Hydrogen bonds are shown as dashed lines and water molecules are depicted as cyan spheres. DNA interacting residues of CUT and HOX domains are shown in green and blue, respectively.

The α3 helix mediates the bulk of DNA contacts by the CUT domain while the loops flanking both ends of α3 and beginning of α2 and α4 helices also bind the DNA. The residues Q353, S364, T367, S369, D370 and R373 of α3 helix as well as K376 in the following loop make hydrogen bonds to the DNA backbone (Fig. 1D). Q353, I351 (in the α2 helix and preceding loop, respectively), K382 and G384 (both in α4 helix) also bind the DNA backbone. S364 and Q365 residues located towards the beginning of α3 helix, and D370 in the same helix are the only residues in CUT making direct base-specific hydrogen bonds with the DNA whereas D370 also makes a water-mediated base interaction. On the other hand, the HOX residues N476, and an arginine pair, R479 and R480, form base-specific hydrogen bonds whereas R450, T469 and N472 interact with the DNA backbone. Overall, the above structural features in OC2 are mostly consistent with those observed in OC1.

A comparison of our OC2 and prior OC1 structure in DNA-bound state to the previous nuclear magnetic resonance (NMR)-based apo-structure of OC1 shows (*24*) that the helix α3 of CUT undergoes a major reorganization upon binding DNA (fig. S1C). In the NMR structure, the beginning of α3 helix, comprising amino acids S364 and Q365, is unstructured while the helix overall is rotated by about 57° compared to that in the DNA-bound form. Intriguingly, the helix α5 of CUT in the DNA-bound structure, although positioned away from the DNA, appears unstructured in the apo-structure. With respect to HOX, the apo- and DNA-bound forms do not show much difference except for the C-terminal stretch beyond helix α8 not being visible in the apo-structure, showing unstructured and flexible nature of this region when not bound to DNA (fig. S1D).

### An OC-specific arginine pair (RR motif) enables unique DNA interaction by the OC2 HOX domain

#### Differences between OC2-PEG10 and OC1-TTR complex structures

Based on our OC2-*PEG10* and previous OC1-*TTR* structures, the bulk of the interactions including base-specific ones, occur through an inner 8 nucleotides of bound DNA (Figs. 1D and Fig. 2, A and B). A comparison of this core region of *TTR* and *PEG10* DNA shows differences mainly at two nucleotide positions – (i) at position 5, a T-A base pair in *TTR* is replaced by a C-G base pair in *PEG10* and (ii) at position 8, a C-G base pair in *TTR* is replaced by a T-A base pair in *PEG10* (Fig. 2, A and B).

**Fig. 2.**
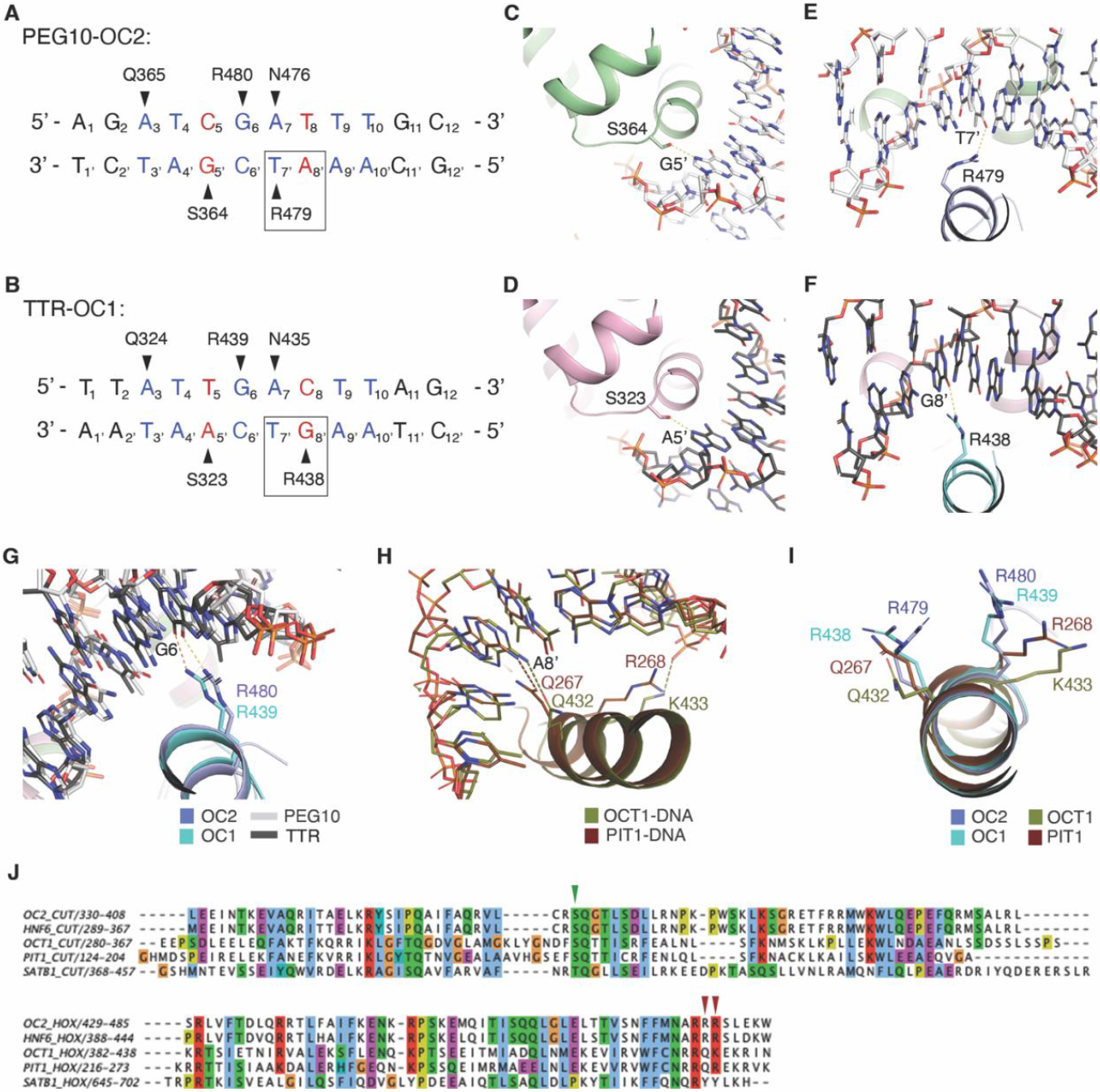
The RR motif in the HOX domain of OC2 shows unique DNA interactions. (**A**) *PEG10* and (**B**) *TTR* DNA sequences and corresponding conserved base-specific interactions with OC2 and OC1, respectively. The core sequence is shown in blue except the bases at which *PEG10* and *TTR* vary, that are in red. Black triangles depict interaction sites. The difference in interaction of the first arginine (OC2 R479 or OC1 R438) of RR pair is shown with a black rectangular outline. Interaction of (**C**) OC2 S364, (**D**) equivalent OC1 S323, and (**E**) OC2 R479 and (**F**) equivalent OC1 R438, to DNA. Hydrogen bonds in panels C to F are shown as yellow dashed lines. (**G**) Interaction of OC2 R480 and equivalent OC1 R439 (yellow and orange dashed lines, respectively). (**H**) Interactions of the POU residues corresponding to the OC arginine pair [PDB 1E3O (OCT1) and 1AU7 (PIT1)]. Hydrogen bonds shown in the same color as respective proteins. (**I**) The relative orientations of the OC arginine pair and corresponding OCT1 and PIT1 residues. (J) Structure-based sequence alignment of CUT (top) and HOX (bottom) domains of OC2, OC1, OCT1, PIT1 and SATB1. The amino acid ranges are indicated. Positions of the conserved serine and the arginine pair of OC2 are shown with green and red arrows, respectively.

The adenine of (5)T-A(5’) base-pair in *TTR* and guanine of (5)C-G(5’) base-pair in *PEG10*, both being purines, form hydrogen bonds through the imidazole N^7^ with a conserved serine side chain oxygen present in both OC1 (S323) and OC2 (S364) (Fig. 2, C and D). This serine is conserved not only within OC family but also in CUT domains of POU members, which show relatively lower sequence similarity to the OC family (Fig. 2J, top). In another related family, SATB, that contains a pair of CUT domains (and a single HOX domain) with even lower sequence conservation to the OC family than POUs, the equivalent residue is a threonine, further showing conservation at this position.

At position 8, the carbonyl oxygen of guanine of (8)C-G(8’) base pair in *TTR* forms a hydrogen bond with a side chain amine of an arginine (R438) located in the helix α8 of OC1 HOX domain (Fig. 2, B and F). However, in the *PEG10* sequence, the nucleotide at position 8’, being an adenine, lacks the carbonyl oxygen needed to form a hydrogen bond to the equivalent side chain of arginine, R479, of OC2. Therefore, the side chain of R479 in OC2 reorients to form a hydrogen bond with the carbonyl oxygen at C^4^ of the preceding thymine (T7’) of *PEG10* DNA (Fig. 2, A and E). Notably, this arginine is part of the arginine pair (RR motif), conserved in the OC HOX domain, that mediate base-specific interactions (as mentioned in section 1; Fig. 2J, bottom). Upon comparison with other common OC recognized promoter sequences, including *HNF-3β, HNF-4, PEPCK and PFK-2GRU* (*8*), we found that the base-pair at position 8 to be variable in these promoter sequences (fig. S2A). This suggests a general sequence variability of OC targeted gene promoters at this position and that the conserved arginine allows OC transcription factors to adapt to this variation.

#### Comparison of the OC RR motif to corresponding POU residues

Having analyzed the binding of the first arginine, as described above, we next examined interaction of the second arginine of the pair (R480 and R439 in OC2 and OC1, respectively) to DNA. R480 in OC2 (and the equivalent R439 in OC1) forms a hydrogen bond with a guanine base (G6) (Fig. 2G). The guanine at this position is, in fact, conserved in the related promoter recognition sequences bound by the OC transcription factors mentioned above (fig. S2A).

The RR motif (R479/R480), although conserved within OC family, is unique relative to POU and SATB members. In POU homeodomains, this motif is made of a glutamine followed by a lysine or arginine (QK/R) whereas in that of SATB it is made of a tyrosine pair (YY) (Fig. 2J, bottom panel). In POU, the glutamine (Q432 and Q267 in human OCT-1 and PIT-1 proteins) corresponding to OC2 R479, forms a hydrogen bond invariably with an adenine in the cognate promoters (Fig. 2H and fig. S2B). The subsequent residue in POU, which is a lysine (K433 in OCT-1) or an arginine (R268 in PIT-1), corresponding to OC2 R480, does not make base contact but can either bind to DNA backbone phosphate or remains unbound (Fig. 2H and fig. S2B), consequently exhibiting difference in orientation relative to OC2 R480 (or OC1 R439) (Fig. 2I). Notably, the base at position 6 in POU specific promoter sequences is generally a cytosine, unlike the corresponding guanine (G6) in OC recognized sequences, which is bound by the second arginine of the RR motif as mentioned above (fig. S2B).

Taken together, the above analyses indicate that the first arginine of the RR motif, through its ability to reorient, confers a certain flexibility to OC2 and OC1 proteins to adapt to cognate base variability in the promoters of OC target genes. In addition, the second arginine, through its interaction with a conserved guanine (G6) in these promoters, provides sequence selectivity.

Furthermore, these base contacts by the RR motif are unique compared to the interactions mediated by the corresponding QK/R stretch found in the POU homeodomains. These observations lead us to propose that this arginine pair is a novel DNA base-pair interacting motif in the HOX domain of OC2.

### The HOX domain thermodynamically stabilizes OC2 on DNA

To understand the mechanism of OC2-DNA interaction further, we carried out thermodynamic analysis of complex formation using isothermal titration calorimetry (ITC). OC2 bound to *PEG10* DNA with a binding affinity (KD) of 6 nM and an associated free energy (ΔG) of -11 kcal/mol. The binding is characterized by a relatively large and favorable enthalpy change (ΔH = -15.3 kcal/mol) and an unfavorable entropy (−TΔS = 4.1 kcal/mol) (Fig. 3, A, D and E).

**Fig. 3.**
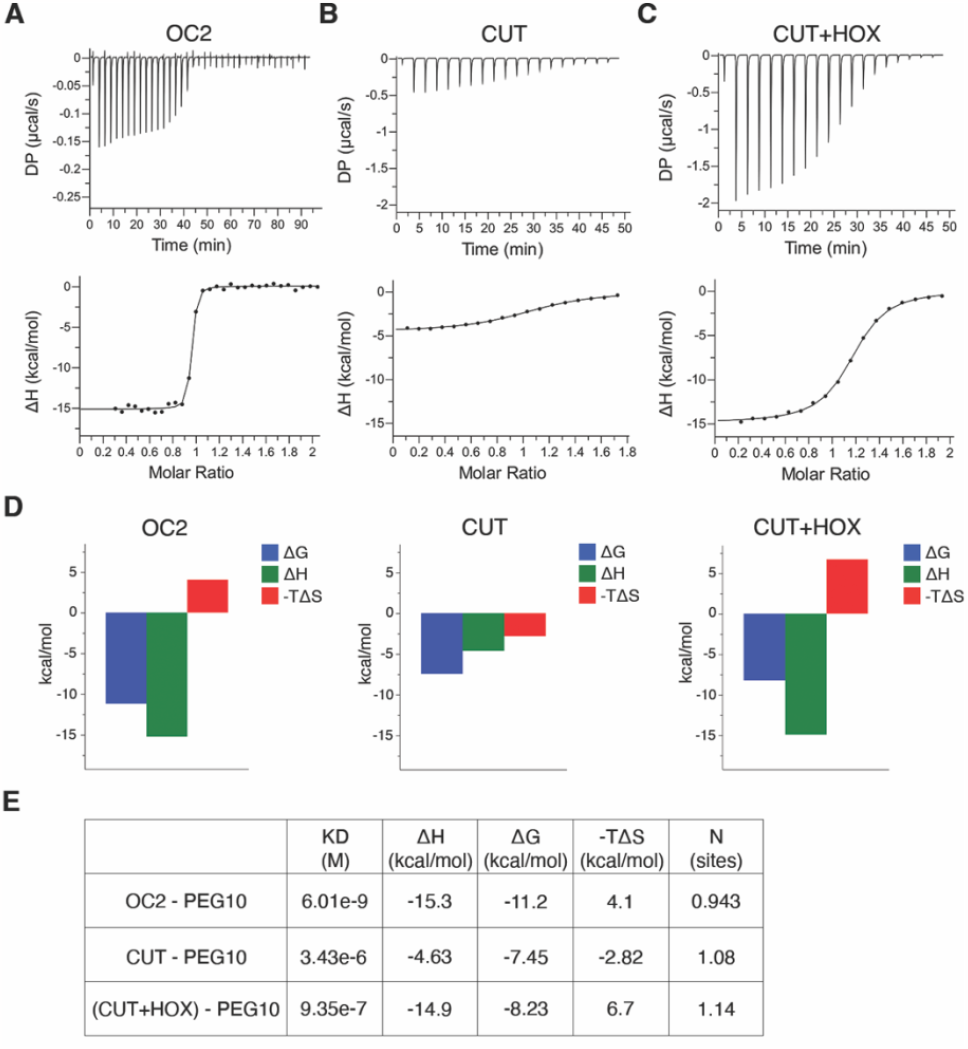
The HOX domain drives stable association of OC2 to DNA. ITC binding analysis of (**A**) intact OC2, (**B**) CUT domain and (**C**) CUT and HOX domains (as separate polypeptides), to *PEG10* DNA. The raw heats (differential power, DP) in microcalories per second for each injection are shown on top and binding isotherms are shown in the bottom. (**D**) Respective thermodynamic signature plots are shown with ΔG in blue, ΔH in green and -TΔS in red. (**E**) KD, ΔG, ΔH and -TΔS values for each reaction are tabulated.

The favorable enthalpy change suggests stable hydrogen bonds and van der Waals interactions being formed while the unfavorable entropy signifies loss of conformational freedom during complex formation. Furthermore, the large favorable enthalpy change compensates for unfavorable entropy resulting in an enthalpically driven interaction. These parameters suggest rearrangements in OC2 upon binding to *PEG10*. Notably, a similar thermodynamic pattern was observed for the interaction between human immunodeficiency virus (HIV) gp120 and CD4 (*25, 26*). HIV gp120 undergoes extensive CD4-induced conformational rearrangements resulting in exceptionally high favorable enthalpy and unfavorable entropy changes during gp120-CD4 complex formation (ΔH = -63 kcal/mol and -TΔS = 51.2 kcal/mol). In the case of the OC2-*PEG10* complex, lower magnitudes of these parameters show relatively smaller regions of OC2 undergoing changes compared to gp120. These conclusions are further supported by the conformational changes observed in the CUT domain, especially in the α3 helix, upon binding to DNA (fig. S1D).

We next sought to understand the roles of the individual CUT and HOX domains towards interaction with *PEG10*. For this, we expressed the CUT and HOX domains separately. The CUT domain bound *PEG10* DNA with nearly 500-fold weaker affinity showing a KD of about 3.4 μM and consequently a significantly lower associated ΔG (−7.4 kcal/mol). Importantly, compared to intact OC2, we observed a markedly lower enthalpy change (ΔH = -4.6 kcal/mol), and strikingly the binding showed a favorable entropy (−TΔS = -2.8 kcal/mol) (Fig. 3B, D and E). These values show a smaller enthalpic and a relatively significant entropic contribution to the overall DNA binding by CUT, indicating a distinct thermodynamic pattern compared to that observed with the intact OC2. We next tested binding of the OC2 HOX domain alone to *PEG10* DNA but observed no binding in this case. We also did not observe any direct binding between CUT and HOX polypeptides in the absence of DNA (fig. S3, A and B).

We then tested whether the presence of HOX, in addition to CUT, but as separate polypeptides, can recapitulate binding of intact OC2 to DNA. We observed a marginal improvement in DNA binding affinity (KD = 0.9 μM) suggesting the covalent linkage provided by an intact linker is needed for optimal binding affinity. Remarkably though, compared to CUT alone, we observed a marked increase in enthalpy change (ΔH = -14.9 kcal/mol) and an unfavorable entropy (−TΔS = 6.7 kcal/mol) (Fig. 3, C to E), a pattern that is similar to one observed with the intact OC2. This marked thermodynamic shift cannot be explained based on an additive effect caused by the HOX based interactions. Further, given the conformational changes associated with this thermodynamic signature, as described earlier, and absence of physical interaction between CUT and HOX, the above observations rather suggest the rearrangements in CUT being induced allosterically by HOX leading to a stable CUT-HOX-DNA ternary complex.

The above observations suggest that CUT initiates DNA-binding, inducing conformational changes in the DNA that facilitates interaction of HOX to the DNA. The DNA-bound HOX allosterically induces rearrangements in CUT, to drive an enthalpy driven and stable DNA interaction of OC2. This data is therefore further indicative of a two-step DNA binding mechanism by OC2 where an initial weaker or transient interaction leads to a subsequent stable complex.

### S364/Q365 and N476 mediated base-interactions are necessary for correct DNA-bound conformation of OC2

An understanding of the role of base-specific interactions in overall DNA binding by OC, and CUT-HOX transcription factors in general, is unclear. Therefore, using the above thermodynamics insights as a template, we sought to understand the contribution of the conserved base-specific interactions towards the DNA binding of OC2. We introduced relevant alanine mutations in both CUT and HOX domains, in context of the intact OC2 DNA binding module. As discussed above, S364 and Q365 residues in the CUT and N476 in the HOX of OC2, form direct base-specific hydrogen bonds with the DNA. These residues are also evolutionarily conserved, across OC, POU and SATB families. In addition, the S364 and particularly Q365 are also conserved in the phage repressors (fig. S4A). Accordingly, we generated two mutants, the first with S364A and Q365A mutations (OC2SQ; double mutant) and the second with N476A (OC2N) mutation. In addition, we also mutated the R479 and R480 (RR motif) (OC2RR; double mutant).

We performed ITC-based DNA binding experiments with these three mutants and compared their thermodynamic parameters with that of wild-type OC2. Both OC2SQ and OC2N mutants bound weaker to *PEG10* DNA with a KD of 46 nM and 61 nM respectively. Importantly, compared to wild-type OC2 (Fig. 3A), both mutants showed significant reduction in the respective enthalpy changes, by almost 30% (ΔH ∼ -10 kcal/mol), while the entropy was significantly more favorable (−TΔS ∼0 kcal/mol) in both cases (Fig. 4, A, B, D and E). These enthalpy and entropy values indicate weaker DNA binding and a disordered complex relative to wild-type OC2. Further, such a large shift in ΔH and -TΔS cannot be solely accounted for by the localized loss of a few hydrogen bonds and suggest that these mutants, lacking proper DNA contacts, are unable to attain the right conformation upon binding to DNA. Notably, the mutated residues S364 and Q365 in OC2SQ are part of α3 helix that undergo structuring upon binding DNA. The observed thermodynamic changes therefore suggest that the base-interactions by S364/Q365 and N476 are essential for the conformational rearrangements in OC2. To test this further, we attempted to crystallize both OC2SQ and OC2N mutants with DNA. However, we failed to obtain any crystals of OC2N while with OC2SQ, we were only able to obtain poor quality crystals that were irreproducible, which might be indicative of the conformational variability and/or disorder in the respective complexes.

**Fig. 4.**
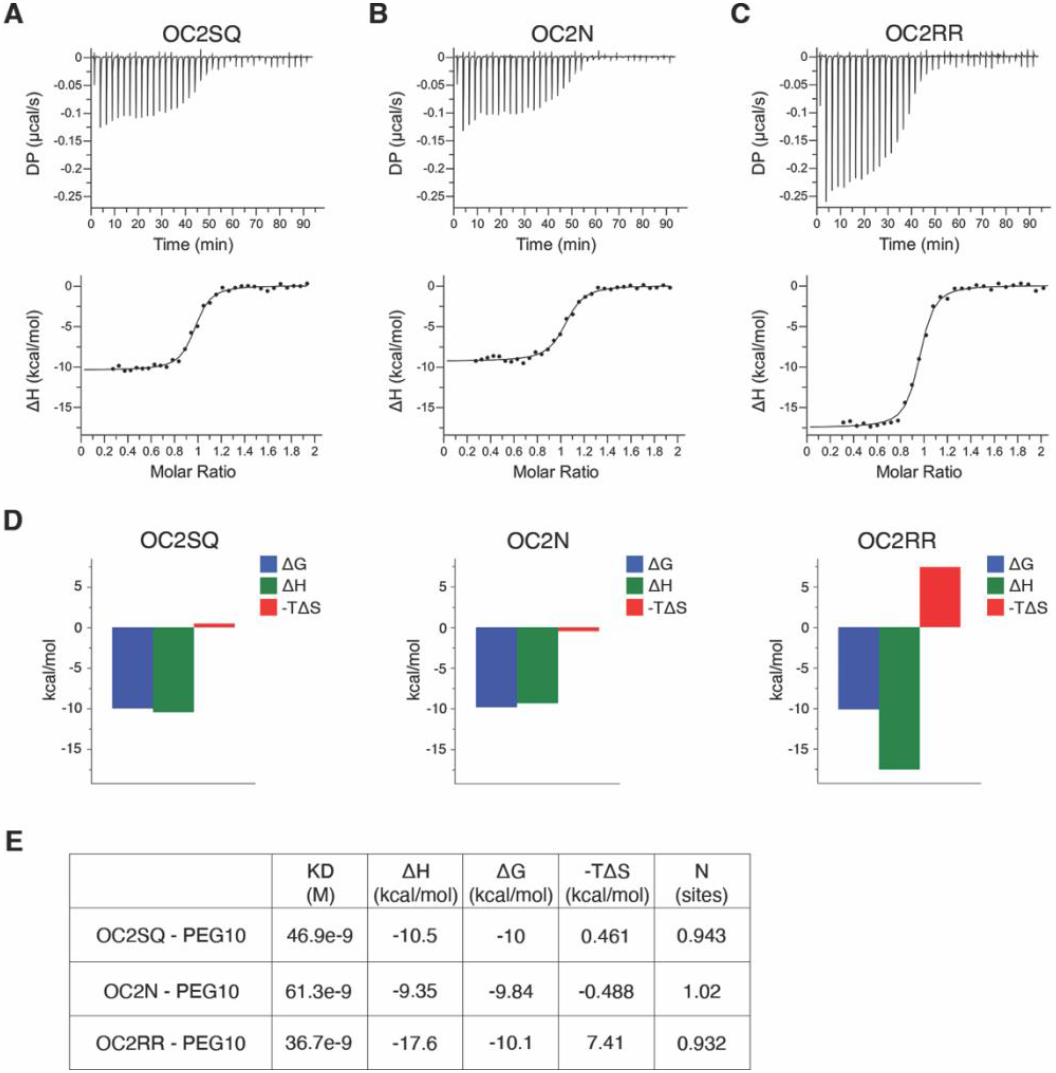
DNA binding thermodynamics of OC2 mutants. ITC binding analysis of (**A**) OC2SQ, (**B**) OC2N and (**C**) OC2RR, to *PEG10* DNA. The raw heats (differential power, DP) in microcalories per second for each injection are shown on top and binding isotherms are shown in the bottom. (**D**) Respective thermodynamic signature plots are shown with ΔG in blue, ΔH in green and -TΔS in red. (**E**) Magnitudes of KD, ΔG, ΔH and -TΔS for each reaction is tabulated. The concentrations of respective mutant proteins and *PEG10* DNA used in each experiment are also shown in the table.

Next, we tested DNA binding by the OC2RR mutant and observed a similarly weaker binding (KD = 36 nM). Intriguingly, in contrast to the other two mutants, OC2RR neither showed a decrease in the enthalpy change (ΔH = -17.6 kcal/mol) nor a favorable change in entropy (−TΔS = 7.4 kcal/mol) (Fig. 4C to E) compared to the OC2-DNA complex. These values indicate a similar conformational state of this mutant in the DNA bound state like that of the wild-type OC2. To further examine this, we crystallized OC2RR with the same *PEG10* DNA and solved the structure at 2.9 Å resolution (table S1). As indicated by the thermodynamic parameters above, this structure mostly resembles that of wild-type OC2-DNA complex.

However, we could not model few additional linker-flanking and the last three C-terminal residues located almost immediately after the RR mutation site, due to disorder (fig. S4B; methods). In addition, surprisingly, we could place only three water molecules in this structure. The reason for this disorder in solvent content is unclear but suggests the ‘RR’ mutations affect solvent stability. These results indicate base-interactions of this arginine pair stabilize the C-terminal of the protein and the overall complex.

Taken together, these data suggest base-specific interactions by S364/Q365, N476, and R479/R480 are needed for optimal DNA binding affinity. However, respective interactions by the evolutionarily conserved S364/Q365 and N476 are essential for accurate OC2 conformation required for favorable DNA binding energetics and are therefore mechanistically separable from that of OC-specific R479/R480.

### Base interactions by OC2, including the RR motif, are essential for optimal DNA binding and transcriptional activity

To further understand the interaction, we studied DNA binding kinetics of the wild-type and mutant OC2 proteins using biolayer interferometry (BLI). We observed that the association of wild-type OC2 to the DNA follows a sigmoidal curve (Fig. 5A and fig. S5A) and the data could be fitted with a 1:2 binding model (see methods; Fig. S5A to D). The association starts (ka1) at a slower rate with eventual faster association (ka2) (Fig. 5E) that manifests as a sigmoid curve.

**Fig. 5.**
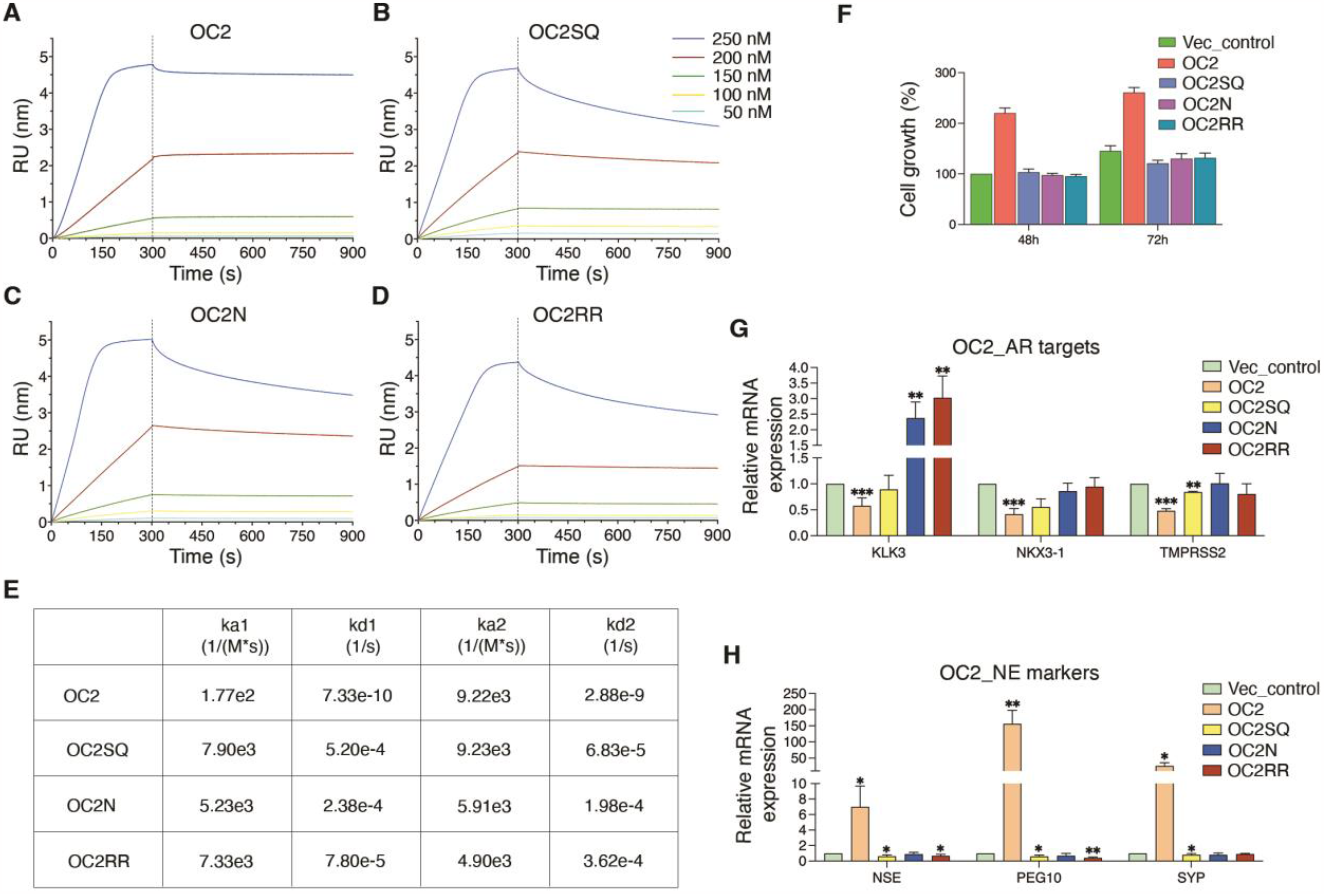
Base-specific interactions by OC2 are needed for DNA binding cooperativity and are functionally relevant in a PC model. Kinetics of *PEG10* DNA binding by (**A**) OC2, (**B**) OC2SQ, (**C**) OC2N and (**D**) OC2RR. All proteins were titrated at concentrations 0, 50, 100, 150, 200 and 250 nM, as shown. The association and dissociation phases are separated by a dotted line. (**E**) The respective rates of association (ka1 and ka2) and dissociation (kd1 and kd2) are tabulated. (**F**) Plot showing the proliferation of LNCaP cells upon stable expression of ectopic wild-type and mutant OC2 compared to cells with endogenous OC2 (vector control) at 48 and 72 hours. Relative mRNA levels of (**G**) AR target genes *KLK3, NKX3-1* and *TMPRSS2* and (**H**) NEPC marker genes *NSE, PEG10* and *SYP*, upon stable overexpression of ectopic wild-type and mutant OC2 compared to cells with endogenous OC2 (vector control). Two-sample t-test was used for statistical analysis. P-values for mRNA levels of genes expressed in cells overexpressing wild-type and mutant OC2 against endogenous OC2 are shown (* P≤0.05, **P≤0.01, ***P≤0.001).

This pattern is indicative of the cooperative nature of the association, like previously reported binding of bacterial protein ParA to DNA (*27*).

We next studied binding kinetics of the three mutants (OC2SQ, OC2N and OC2RR) to DNA. We observed an absence of the initial sigmoidal association in all the mutants (Fig. 5, B to D). This is reflected in the increase in the first or initial association rate, ka1 (Fig. 5E), and all the mutants gained an order of magnitude (30 to 45-fold increase) in ka1 although their second association rate (ka2) remained similar, resulting in a smoothening of the sigmoid association (towards hyperbolic) in case of the mutants. Furthermore, all three mutants dissociated significantly faster (4 to 5 orders of magnitude) relative to the wild-type OC2 (Fig. 5E). The non-sigmoidal association pattern shows a loss of DNA binding cooperativity in all three mutants.

The kinetics further suggest that the base-specific interactions by wild-type OC2 causes a slower association as well as dissociation, thereby stabilizing the complex. The kinetics data further show that the OC2RR mutant, despite exhibiting contrasting DNA binding thermodynamics than OC2SQ and OC2N mutants, is also defective in DNA binding.

We further evaluated functional relevance of these base-specific interactions in prostate cancer context. Our earlier work and that by Guo et al. (*12, 13*) showed that overexpression of OC2 leads to AR axis suppression and development of NEPC characteristics (lineage plasticity) in LNCaP cells, an androgen receptor (AR) dependent prostate cancer model characterized by mild endogenous OC2 expression. Upon constitutive overexpression of the OC2 SQ, N or RR mutant, instead of the wild-type OC2 protein, we found that the proliferation of the cells was reduced to that observed at endogenous OC2 levels (Fig. 5F). We analyzed mRNA levels of three PC relevant AR target genes *KLK3, NKX3-1*, and *TMPRSS2*, in cells expressing either OC2SQ, OC2N or OC2RR mutants. Unlike the wild-type OC2, none of the three OC2 mutants suppressed these AR targets (Fig. 5G). Lastly, we tested expression of three NE differentiation markers *NSE, PEG10* and *SYP*, that are upregulated by OC2. Consistently, none of the mutants upregulated these genes (Fig. 5H). These results validate the interactions we have identified and characterized biochemically are necessary for OC2 transcriptional activity in the prostate cancer model tested.

## Discussion

We report here an analysis of the DNA binding mechanism of OC2, a developmentally important transcription factor and a driver of treatment resistant prostate cancer. We show that the HOX domain is critical in driving an energetically favorable OC2-DNA complex by allosterically inducing rearrangements in the CUT domain. This implies a two-step mechanism of cooperative DNA binding by OC2 wherein initial contacts to DNA are made by CUT followed by binding of HOX that thermodynamically stabilizes OC2 onto DNA. Remarkably, the energetics pattern of the OC2-DNA complex is reminiscent of HIV gp120-CD4 interaction that has been harnessed for development of potent antivirals (*28, 29*). In parallel structural studies, we identified a unique DNA base-interacting arginine pair in the HOX domain of OC2, which we call the ‘RR motif’. This amino acid pair is unique to the OC family compared to POU and SATB. In addition, the first arginine interacts distinctly to DNA in respective OC2-*PEG10* and OC1-*TTR* complexes, suggesting a mechanism to tolerate specific alterations in OC promoter sequences with implications on the redundant transcriptional activation by OC paralogs (*8, 30*). These findings together demonstrate the HOX domain to be a key regulatory element for OC2-DNA binding.

Probing the mechanism further, we found the base-specific contacts by S364/Q365 in CUT and N476 in HOX, residues conserved across OC, POU and SATB families, to be essential determinants for an energetically favorable and therefore, conformationally correct OC2-DNA complex. Nonetheless, base interactions by OC specific R479/R480 (RR motif), apart from S364/Q365 and N476, are needed for optimal DNA binding affinity, kinetics, and cooperativity. Notably, in a prostate cancer model, we show these interactions to be functionally relevant in terms of transcriptional activity and disease progression. Collectively, these findings demonstrate that the respective DNA interactions by the evolutionarily conserved amino acids S364/Q365 and N476 ensure the basic functional framework while family-specific elements in the HOX domain, like the RR motif in OC, provide additional mechanistic properties. Overall, this is the first report attributing a structural role to DNA base-specific interactions by CUT-HOX proteins.

In conclusion, we propose that the HOX domain with its crucial thermodynamic contribution and unique RR motif, regulates stable interaction between OC2 and DNA. Our integrative approach reveals unprecedented molecular details of DNA binding by OC2 with broad mechanistic implications for CUT and related POU family transcription factors, and that present potential new therapeutic opportunities for intervention.

## Acknowledgments

The authors thank staff at X-ray and EM structure determination core at UCLA for granting access to Octet RED 96 instrument.

## Author contributions

R.M., M.R.F. and A.C. designed the research; R.M. and M.R.F. supervised the work. A.C. performed protein purifications, crystallization, structure analysis, mutant design, and kinetics studies. M.K. and A.C. carried out structure determination and refinement. B.G. performed ITC experiments. M.K., M.R.H., and B.G. helped with protein purifications. C.Q. generated stable cell-lines, performed cell-proliferation and gene expression analyses. A.C. wrote and edited the original draft with critical reading by M.K., B.G., R.M. and M.R.F. and inputs from all authors. M.R.F. and R.M. acquired the funding.

## Data and materials availability

Contact R.M. and M.R.F. for reagent requests. The PDB files of the structures with accession codes 8T0F (OC2-PEG10) and 8T11 (OC2RR-PEG10) are available at www.rcsb.org.

## Supplementary Materials

Materials and Methods

Figs. S1 to S6

Table S1

References (*31-40*)

