## Supplemental material for "The homeodomain drives favorable DNA binding energetics of prostate cancer target ONECUT2"

**The PDF file includes:**

Materials and Methods
Figs. S1 to S6
Table S1
References (*31-40*)

### Materials and Methods

Protein expression and purification

The human OC2 DNA binding region spanning residues 330-485 (OC2) was cloned into pET-His6-TEV-LIC expression plasmid (Addgene Plasmid #29653). The protein was expressed in *Escherichia coli* (*E. coli*) BL21(DE3) cells. The cells were grown at 37 °C to an optical density (OD) of 0.8 in Terrific Broth (TB) media and induced with 0.5 mM isopropyl β- d-1-thiogalactopyranoside (IPTG) at 18 °C overnight. Cells were lysed by sonication in buffer containing 50 mM Tris pH 7.5, 500 mM NaCl, 20 mM Imidazole, 10% Glycerol and 5 mM β-mercaptoethanol (β-ME) (Buffer A). Cell debris were removed by centrifugation at 43,600 x*g* and cleared lysate was passed through nickel-nitrilotriacetic acid (Ni-NTA) resin (Qiagen). The protein was eluted in Buffer A supplemented with 500 mM Imidazole. The His-tag was removed by incubating Ni-NTA eluate with Tobacco Etch Virus (TEV) protease at 4 °C overnight. The sample was diluted to reduce NaCl and imidazole concentrations to 50 mM each and passed through Ni-NTA resin again to remove the cleaved His-tag and TEV (also His-tagged). The protein was then loaded on a 5 mL HiTrap SP (Cytiva) cation exchange column, equilibrated in buffer containing 25 mM HEPES pH 7.4, 50 mM NaCl, 10% Glycerol and 1 mM DTT and eluted with a linear gradient of 50 mM to 1 M NaCl. The fractions containing OC2 were concentrated and loaded onto a Superdex S75 gel filtration column (Cytiva) equilibrated with buffer containing 25 mM HEPES pH 7.5, 250 mM NaCl and 1 mM DTT. The purified protein was aliquoted, flash-frozen in liquid nitrogen and stored at -80 °C. All the mutants were also prepared using the same protocol. The final purified wild-type and mutant OC2 showed similar SDS-PAGE and gel-filtration elution profiles (fig. S6). The OC2 residues 317-417, containing the CUT domain, and residues 420-490, containing the HOX domain were cloned into pET-His6-MBP-TEV-LIC expression plasmid (Addgene Plasmid #29656). Both proteins were purified using the same protocol described above for the intact OC2 protein.

Site-directed mutagenesis

Mutations in OC2 DNA binding region (residues 330-485) for purified protein-based studies were introduced by site-directed mutagenesis using Pfu Turbo (Agilent) DNA polymerase. The PCR product was treated with DpnI (NEB) enzyme at 37 °C for 1 hour and transformed into Top10 *E. coli* cells. Mutagenesis in full length *OC2* for the cell-based assays were performed using Quick Change II XL site-directed mutagenesis kit (Agilent) according to manufacturer's protocol. Mutations were confirmed by DNA sequencing.

Crystallization, data collection and structure determination

OC2 DNA binding site was originally mapped to 14 base pairs within *PEG10* promoter sequence (*12*), so, we initially attempted to co-crystallize OC2 with the corresponding 14 mer DNA duplex (fig. S1E). However, this 14 mer DNA yielded crystals that were difficult to reproduce. Changing the DNA to a 12 mer duplex, lacking one base pair from each terminus in comparison to the 14 mer sequence, resulted in crystals that formed more readily.

DNA oligos purchased from IDT were annealed for crystallization and duplex DNA formed was mixed with protein in 1:1.4 ratio (protein to duplex DNA). Crystallization was set-up at 18 °C by hanging drop vapor diffusion method. OC2-*PEG10* complex crystals were obtained in the condition 0.04 M $KH_2PO_4$, 16 % PEG 8K and 20 % Glycerol while OC2RR-*PEG10* complex crystals appeared in the condition 10 % PEG 1K and 7.5 % PEG 8K. Data were collected in an in-house Rigaku Micromax 007 HF rotating anode X-ray generator and R-axis IV++ image-plate detector. Data processing was performed with HKL2000 (*31*). Structure determination of OC2-*PEG10* was carried out by molecular replacement method using MolRep (*32*), with the OC1-*TTR* complex structure (PDB 2D5V) as a search model. For the OC2RR-*PEG10* structure solution, OC2-*PEG10* structure was used as a search model. Model building was done with COOT (*33*) while refinement was carried out using REFMAC (*34, 35*) and Phenix Refine (*36*). In both structures, the amino acids 409-428, representing the linker, could not be modeled due to lack of electron density. In addition, the OC2RR-*PEG10* structure also lacked proper electron density for residues 407-408, 429-432 and 483-485. The data collection and refinement statistics are provided in table S1. Structure figures were prepared with PyMol (The PyMOL Molecular Graphics System, Version 2.4 Schrödinger, LLC). All the above crystallographic softwares were used from the SBgrid platform (*37*). Protein-DNA interaction map was prepared with LigPlot+ (*38*).

ITC binding studies

ITC experiments were performed using MicroCal PEAQ-ITC (Malvern Panalytical). Both protein and DNA were dialyzed in 1X phosphate buffered saline (PBS) pH 7.4. For experiments involving intact (wild-type and mutant) OC2, duplex DNA at 100 μM (in syringe) was titrated as 36 injections of 1 μL each against 10 μM protein (in the cell). For experiments involving isolated CUT and HOX domains, five-fold higher concentrations of protein and DNA were used due to lower heats generated. Accordingly, for these experiments, DNA at 500 μM (in the syringe) was titrated as 18 injections of 2 μL each against 50 μM protein (in the cell). The data was processed with MicroCal PEAQ-ITC Analysis software.

Biolayer interferometry (BLI) kinetic studies

Protein-DNA kinetic studies were carried out in Octet RED96 (Sartorius ForteBio). One of the *PEG10* oligos was biotinylated on the 5'-end (IDT) and annealed to the non-biotinylated complimentary oligo. This biotinylated duplex DNA was immobilized on a SADH biosensor (Sartorius) and OC2 (wild-type or mutants) was titrated at 0, 50, 100, 150, 200 and 250 nM concentrations. The assay was performed in 1X phosphate buffered saline (PBS) + 0.5 mM tris [2-carboxyethyl] phosphine (TCEP). The 250 nM curve for all proteins showed a slightly different profile from the rest. Due to this difference, fitting of all the curves using a single binding model was not possible and additionally, the 250 nM curve especially showed poor fitting to any binding model. We therefore omitted the 250 nM curve for calculating the kinetic parameters. Data was fitted with 1:2 model using TraceDrawer software (Ridgeview Instruments).

Protein sequence alignments

Alignments were performed using Clustal Omega (*39*) and edited in Jalview (*40*).

Stable Cell line Generation

LNCaP (#CRL-1740) was obtained from the American Type Culture Collection (ATCC) and authenticated using the Promega PowerPlex 16 system DNA typing (Laragen). Mycoplasma contamination was routinely monitored using the MycoAlert PLUS Mycoplasma Detection Kit (Lonza). The OC2 overexpression construct was generated by cloning the full-length OC2 cDNA

(NM_004852) into the pLenti-C-Myc-DDK-IRES-Puro (Origene PS100069) lentivirus system. Then packing (psPAX2, Addgene #12260), and envelope (pMD2.G, Addgene #12259) plasmids were co-transfected into HEK293T cells to produce lentivirus. Cells were infected with lentivirus supplemented with 10 µg/mL polybrene, then selected by 2ug/mL puromycin to generate the stable overexpression cells. All cell lines were grown in RPMI-1640 media (Gibco) supplemented with 10% FBS and penicillin/streptomycin.

Relative mRNA expression levels of endogenous *OC2* (vector control), wild-type *OC2*, *OC2SQ*, *OC2N* and *OC2RR* in respective stably expressing LNCaP cells are shown in Fig. S5E.

Cell proliferation analysis

All procedures were performed according to the XTT cell viability kit protocol (CST). Seeding was done with 2000 cells/well and grown up to 72 hours, then absorbance at 450nM was measured for further analysis.

RT-qPCR for gene expression analysis

Total RNA from cells was extracted using Qiagen RNeasy Kit (Qiagen) following the manufacturer's instructions. 1µg of total RNA was reverse transcribed to cDNA with iScript cDNA Synthesis Kit (Bio-Rad) following manufacturer's instructions. 2X PowerUp SYBR Green Master Mix (ThermoFisher) was used for cDNA amplification. Assays were performed in triplicates and normalized to β-actin. Graphs were prepared using GraphPad Prism (www.graphpad.com).

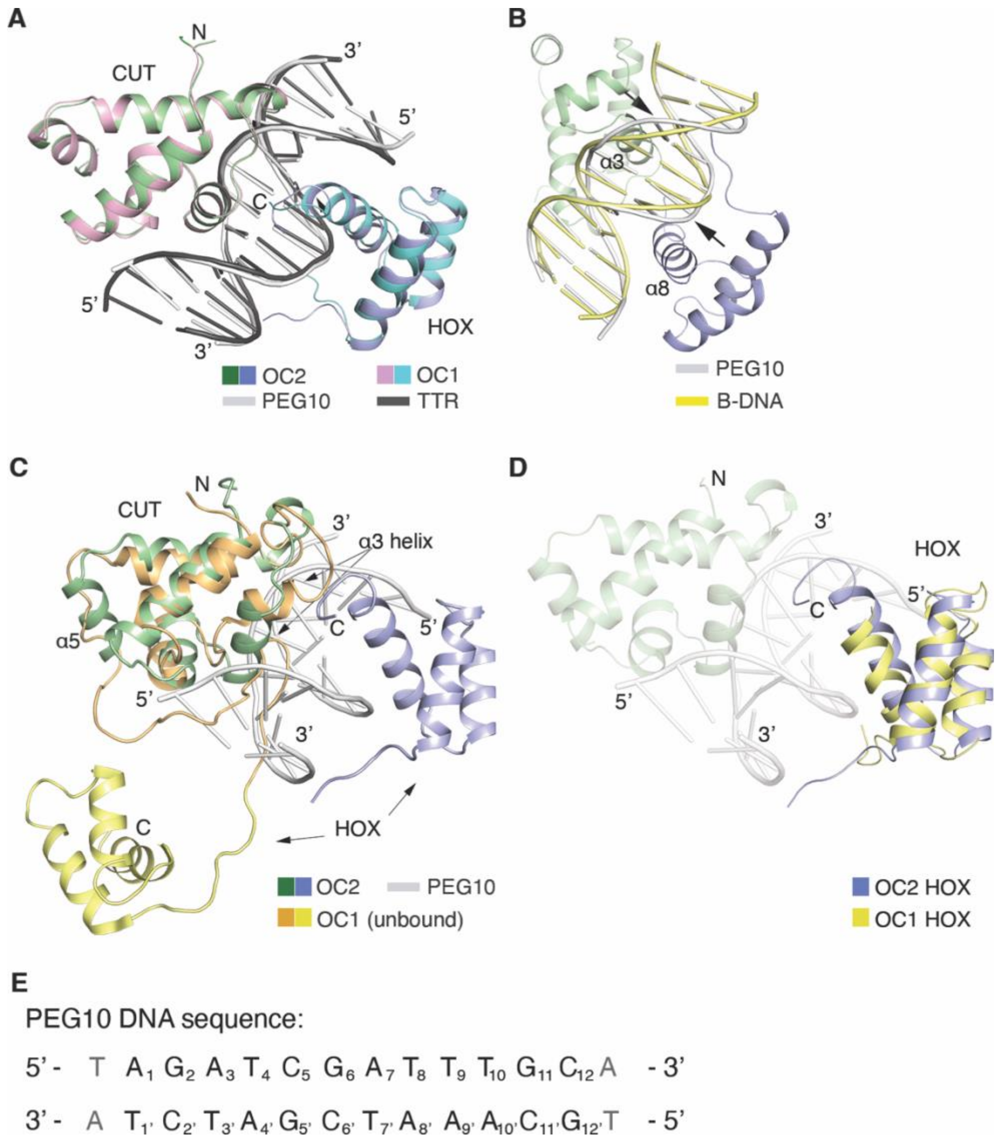


**Fig. S1.**

**Structural analyses of OC2-*PEG10* complex.**
(A) Alignment of OC2-*PEG10* and OC1-*TTR* (PDB 2D5V) structures. The structures align with an RMSD of 0.78 Å. (B) Alignment of OC2-bound PEG10 DNA and B-DNA. The bulge in the major groove is marked with arrows. (C) Relative orientations of OC2 and OC1 in DNA bound and unbound (PDB id 1S7E) (*24*) states, respectively, with the CUT domains from the two structures aligned. The α3 and α5 helices are labeled. (D) Alignment of the HOX domains of the DNA bound OC2 and DNA unbound OC1. (E) The 14mer PEG10 target DNA sequence for OC2 binding is shown. The 12mer sequence used in this study is shown in black, and numbered accordingly, with the excluded base pairs in grey. The N- and C-termini of the structures are labeled as also the 3’ and 5’ of the DNA.

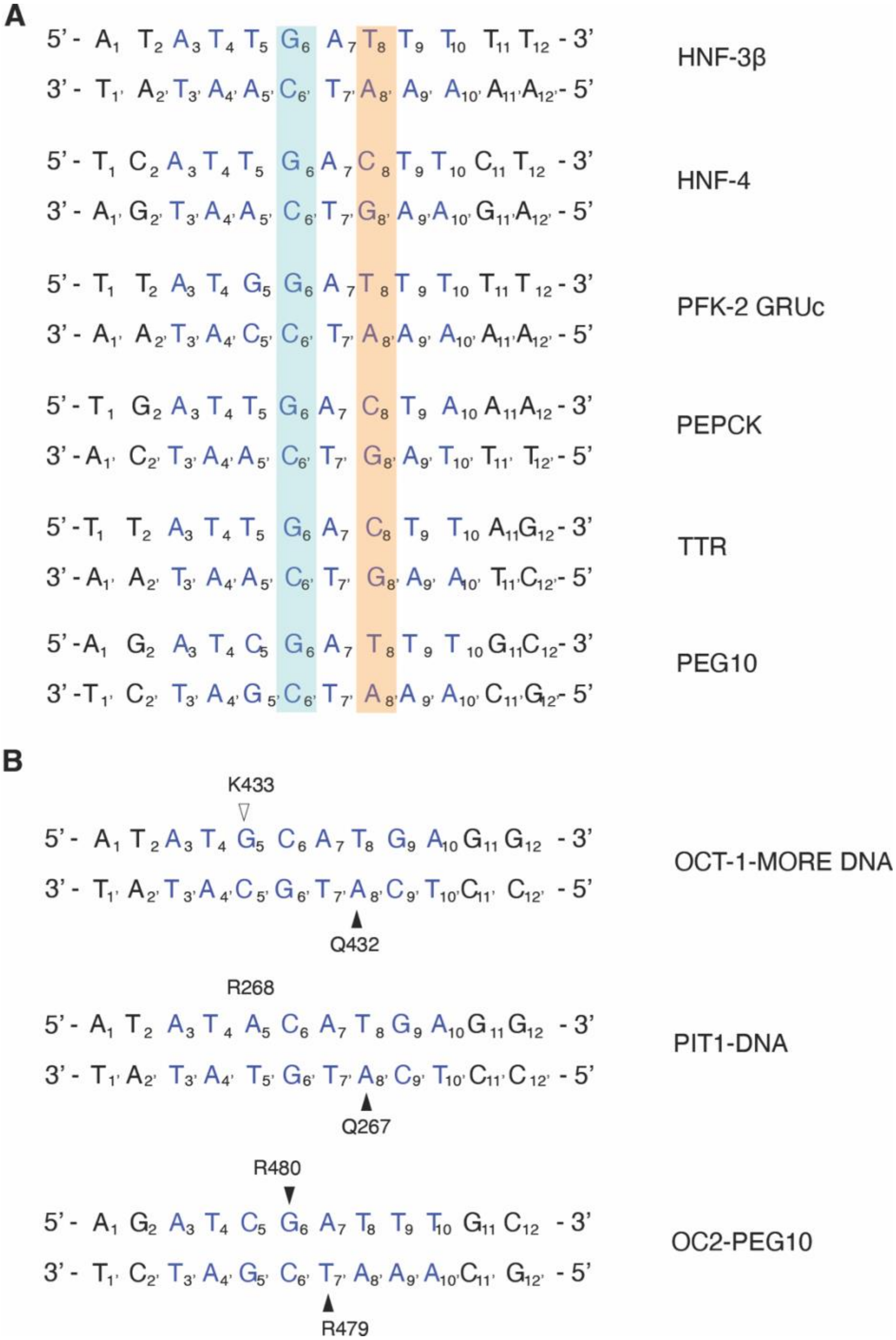


**Fig. S2.**

**OC and POU target promoter sequence analyses.**

(A) Common OC target promoter sequences, *HNF-3β* (hepatocyte nuclear factor-3β, *HNF-4* (hepatocyte nuclear factor-4), *PFK-2 GRUc* (6-phosphofructo-2-kinase glucocorticoid response unit c site), *PEPCK* (phosphoenolpyruvate carboxykinase) and *TTR* (transthyretin) (*8*). The *PEG10* sequence is also shown for reference. Conservation of the guanine base at position 6, bound by OC2 R480 (or OC1 R439), is highlighted in cyan. The variable base at position 8 is highlighted in orange. (B) Sequence of DNA bound to OCT1 (*MORE* DNA; PDB 1E3O) and PIT1 (PDB 1AU7) (as labeled on the right). The OC2 bound *PEG10* sequence is also shown (bottom) for reference. Note the conserved adenines at position 8’ in OCT1 and PIT1 bound DNA that interacts with the conserved glutamine (OCT1 Q432 and PIT1 Q267; shown with solid black triangles) and the difference in position 6 of OC and POU bound DNA sequences. Interaction of OCT1 K433 to DNA backbone carbonyl group is depicted with an open black triangle. PIT1 R268 does not show any interaction in the corresponding structure.

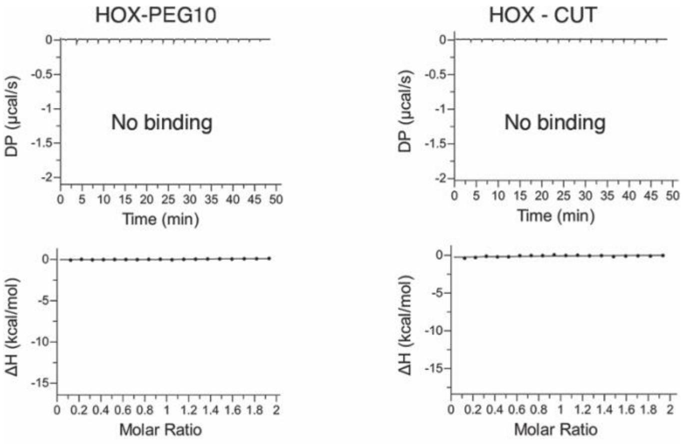


**Fig. S3.**

**ITC binding analysis of (A) HOX-*PEG10* and (B) HOX-CUT.**

The raw heats (differential power, DP) in microcalories per second for each injection are shown on top and binding isotherms are shown in the bottom.

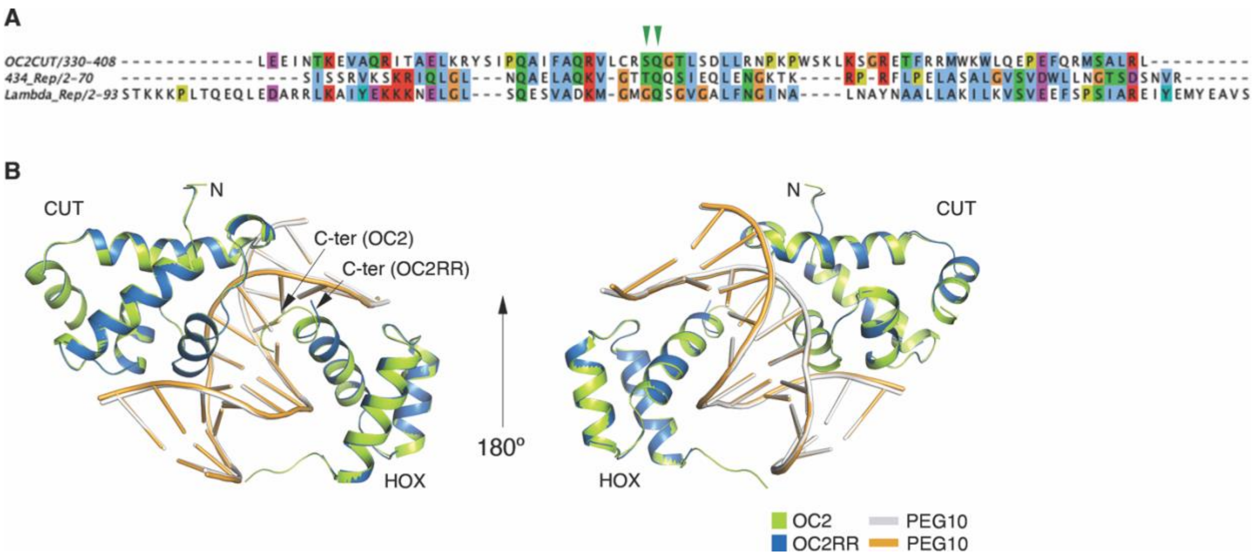


**Fig. S4.**
Structure-based sequence alignment of DNA bound OC2 CUT, 434 phage repressor (PDB 2OR1) (*17*) and Lambda phage repressor (PDB 1LMB) (*20*) domains. Positions of the conserved serine and glutamine of OC2 CUT domain are shown with green triangles. The amino acid ranges for respective sequences are indicated. (B) Structural alignment of OC2 and OC2RR mutant (RMSD=0.296 Å). The CUT and HOX domains are labeled. The N-terminal is also labeled while the C-terminal of both structures are marked with arrows in the left panel.

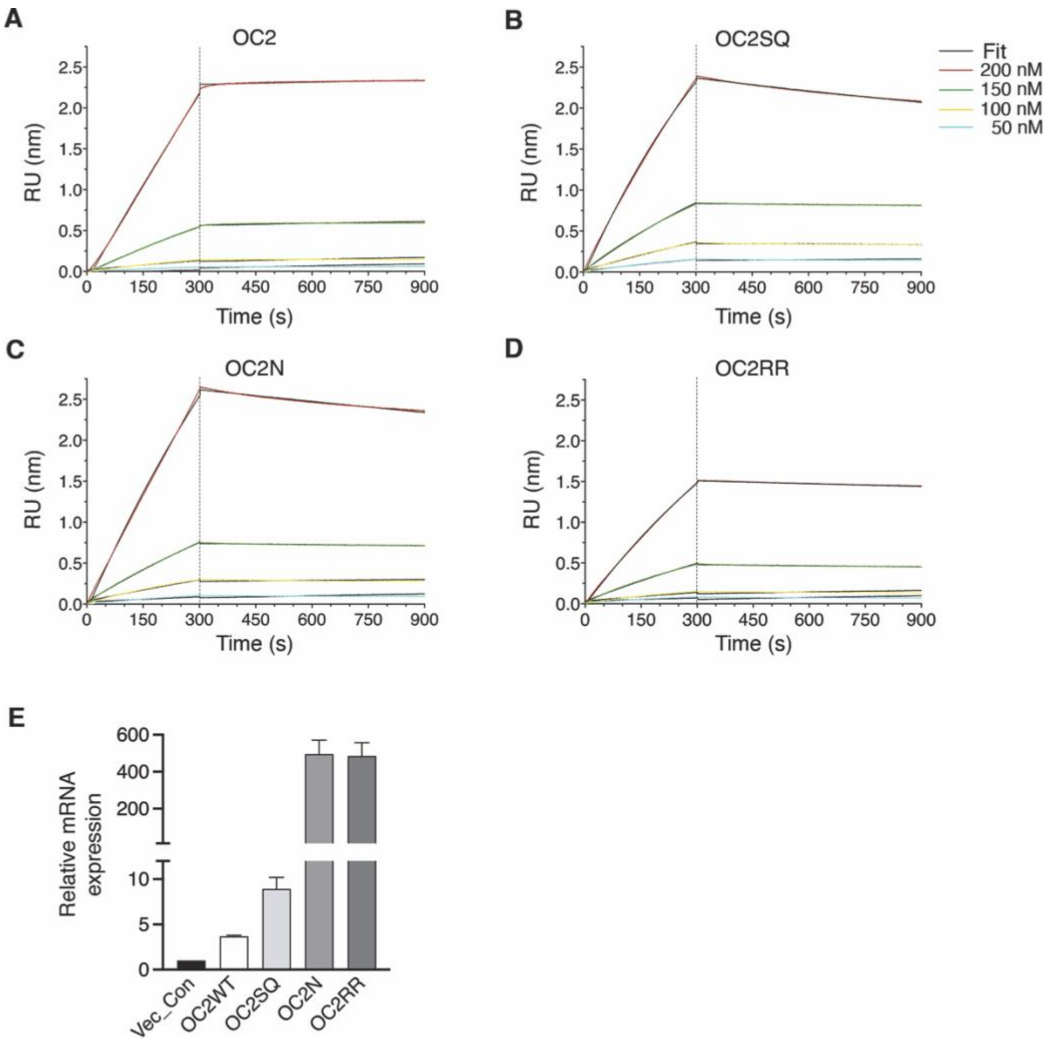


**Fig. S5.**
(A-D) Association and dissociation curves of wild-type and mutant OC2 with respective fits to 1:2 binding model for the 50, 100, 150 and 200 nM curves (see methods). (E) Relative mRNA levels of endogenous *OC2*, *OC2 wild-type*, *OC2SQ*, *OC2N* and *OC2RR*, in respective LNCaP cells.

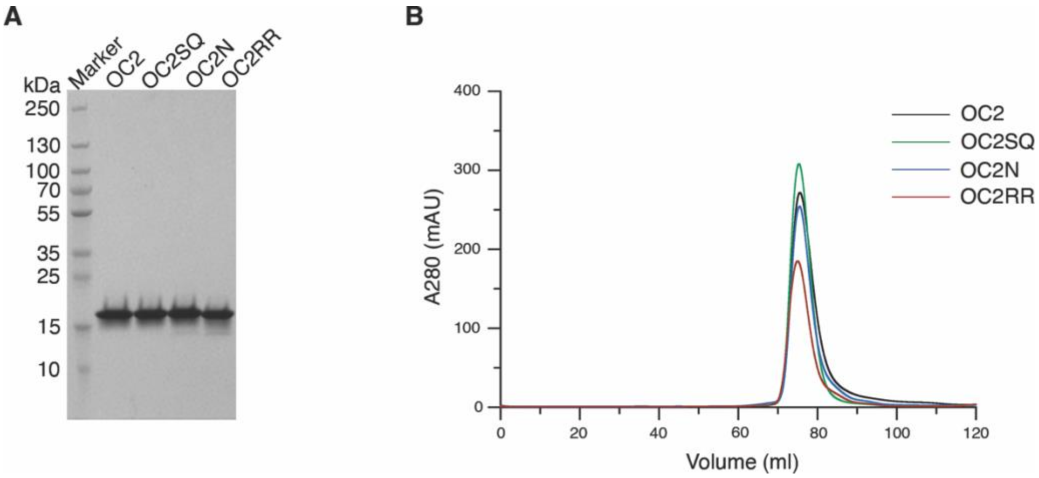


**Fig. S6.**
**Purification profiles of OC2 and its mutants.**
(A) SDS-PAGE gel showing profiles of all purified proteins. The molecular weights of marker bands are shown in kDa. (B) Gel-filtration (Superdex 200 16/600) profiles of all proteins are shown.

**Table S1.**
OC2-*PEG10* and OC2RR-*PEG10* crystallographic data collection and refinement statistics

| Parameter | OC2-*PEG10* | OC2RR-*PEG10* |
|---|---|---|
| PDB code | 8T0F | 8T11 |
| Wavelength (Å) | 1.5418 | 1.5418 |
| Resolution range (Å) | 28.39 - 2.61 (2.703 - 2.61) | 27.87 - 2.91 (3.014 - 2.91) |
| Space group | C 1 2 1 | C 1 2 1 |
| Unit cell parameters | | |
| a, b, c (Å) | 96.229, 78.303, 39.447 | 96.14, 78.154, 39.501 |
| α, β, γ (°) | 90, 110.993, 90 | 90, 111.436, 90 |
| Total reflections | 60782 (6008) | 21153 (2129) |
| Unique reflections | 8316 (818) | 5960 (582) |
| Multiplicity | 7.3 (7.3) | 3.5 (3.7) |
| Completeness (%) | 98.56 (98.20) | 97.38 (95.65) |
| Mean I/sigma(I) | 21.89 (3.06) | 11.81 (1.61) |
| Wilson B-factor ($Å^2$) | 52.34 | 56.44 |
| R-merge | 0.1195 (1.119) | 0.1575 (1.277) |
| R-meas | 0.1286 (1.203) | 0.186 (1.493) |
| CC1/2 | 0.999 (0.712) | 0.994 (0.73) |
| CC* | 1 (0.912) | 0.998 (0.919) |
| Reflections used in refinement | 8269 (818) | 5881 (572) |
| Reflections used for R-free | 431 (51) | 434 (43) |
| R-work | 0.2290 (0.3531) | 0.2511 (0.4423) |
| R-free | 0.2835 (0.4851) | 0.3058 (0.6158) |
| CC (work) | 0.953 (0.729) | 0.940 (0.640) |
| CC (free) | 0.928 (0.553) | 0.902 (0.604) |
| Number of non-hydrogen atoms | | |
| Total | 1668 | 1535 |
| Macromolecules | 1631 | 1532 |
| Ligands | 0 | 0 |
| Solvent | 37 | 3 |
| Protein residues | 135 | 126 |
| RMSD from ideal | | |
| Bond lengths (Å) | 0.004 | 0.003 |
| Bond angles (°) | 0.61 | 0.57 |
| Ramachandran Plot | | |
| Favored (%) | 96.95 | 98.36 |
| Allowed (%) | 3.05 | 1.64 |
| Outliers (%) | 0.00 | 0.00 |
| Rotamer outliers (%) | 0.00 | 0.00 |
| Clashscore | 7.13 | 4.88 |
| Average B-factor ($Å^2$) | | |
| Overall | 55.57 | 45.77 |
| Macromolecules | 60.86 | 48.78 |
| Solvent | 52.08 | 31.28 |

Values in parentheses are for the highest resolution shells
